# Large-scale HLA immunopeptidome and interactome profiling in microglia

**DOI:** 10.1101/2025.04.23.650327

**Authors:** Sydney Klaisner, Ying Hao, Alexandra Beilina, Jae-Hyeon Park, Ziyi Li, Michiyo Iba, Cole Tindall, Benjamin Jin, Jacob Epstein, Isabelle Kowal, Caroline B. Pantazis, Nicole Carmiol, Paige Jarreau, Nicole Washecka, Cory Weller, Mike A. Nalls, Kendall Van Keuren-Jensen, Andrew B. Singleton, Mark R. Cookson, Yue Andy Qi

## Abstract

Microglia are immune cells of the brain and act as major antigen presenting cells. Antigen presentation involves the human leukocyte antigen (HLA) complex, which is implicated in genetic risk of multiple neurodegenerative diseases. How HLA affects the function of microglia in the context of neurodegenerative disease remains unclear. Here, we investigated the HLA epitopes and their protein interactome in human induced pluripotent stem cell (iPSC)-derived microglia-like cells (iMGLs) using systematic mass spectrometry (MS)-based immunopeptidomics, whole-cell proteomics, affinity purification, and prediction algorithms. Our results revealed the presence of almost 7,000 peptides presented by HLA class I and II within microglia. We further showed that the immunopeptidome landscapes of iPSCs, iMGLs and interferon-gamma (IFNγ) stimulated iMGLs are all readily distinguishable. Furthermore, HLA interacts with different groups of proteins in iPSCs compared to iMGLs which involve proteins in immune response. Importantly, we detected 25 HLA epitopes derived from 15 genes associated with Alzheimer’s and related dementias such as Tau, PLD3 (Alzheimer’s disease), TDP-43, FUS (Frontotemporal dementia), and PARK7, VPS35 (Lewy Body dementia). We predicted 31 mutant epitopes derived from these ADRD genes that could be presented with strong interaction to HLA molecules. Along with these epitopes, we observed an enrichment of immune-related interaction proteins in microglia treated with IFNγ. These results provide evidence that aggregated and mutated proteins can interact with HLA alleles and be presented on the cell surface by microglia cells. This study sheds light on the antigen presenting and adaptive immunity mechanism within the central nervous system and its possible effects on neurodegenerative diseases.

## Introduction

In the central nervous system (CNS), microglia are innate immune cells that provide support and monitoring for pathogens and foreign proteins [1]. Although the CNS has been thought to be immune privileged, activated microglia express increased levels of HLA antigens and become phagocytic, engulfing many target proteins [1, 2]. Additionally, microglia have shown to interact and modify synapses through “synaptic stripping” affecting functional transmission and circuit function [3]. Neuroinflammation in neurodegenerative disorders, such as Alzheimer’s disease (AD) and frontotemporal lobar dementia (FTLD), is characterized in part by a change in microglia from quiescent to multiple activated states [4, 5]. While microglial plasticity is likely adaptive over an acute time frame, it is thought that chronic activation associated with neurodegenerative disease may result in neuronal damage due to microglia shifting to disease specific states [6]. Microglia also act as antigen-presenting cells (APCs) in the brain. APCs are involved in processing and presenting foreign or self-proteins by loading epitopes of a specific peptide length onto the peptide binding groove present on the human leukocyte antigen (HLA). HLA molecules then present the epitopes to the cell surface [7]. This HLA-epitope complex has specific recognition towards T-cell receptors (TCRs) and is crucial in the development of an adaptive immune response. In the context of neurodegeneration, epitopes developed from neurodegenerative related genes may form an interaction with HLA [8]. Recognition of this epitope by peripheral lymphocytes leads to an influx of immune cells entering the CNS causing progressive neuronal degeneration and neuroinflammation (**Figure S1**).

Mass spectrometry (MS) has been widely used to provide targeted profiling and detection of specific peptides in a relatively unbiased manner. For immunopeptidomics, MS enables characterization of the antigen presenting mechanism and provides experimental detection of HLA epitopes across various diseases [9-11]. Peptide interaction and affinity with a variety of HLA alleles is critical in understanding the activation of the adaptive immune response and have been extensively studied through prediction algorithms [11, 12]. A neural network-based algorithm NetMHCpan, integrates broad MHC coverage with MS peptidome data leading to improved characterization compared to other neural network models [11, 13].

Despite MS-based HLA profiling being mainly applied in various epithelial cells and blood cells for cancer immunotherapy, the landscape of HLA profiling remains unexplored in central nervous system cells. To understand the mechanism of HLA-peptide interactions and its correlation to the etiology of neurodegenerative diseases, we analyzed the HLA immunopeptidome and interactome in human pluripotent stem cell (iPSC)-derived microglia-like cells (iMGLs) by combining affinity purification and MS-based peptidomics. In this study, we conducted HLA typing, immunopeptidome and HLA interactome profiling in the microglia cells that were derived from the KOLF2.1J iPSC line. Additionally, we stimulated microglia with interferon-gamma (IFNγ), a key regulator and activator of microglia and other macrophages, which is known to induce a phenotype with expression of inflammatory cytokines and induces both the innate and adaptive immune responses [14, 15]. We showed distinctive HLA epitope signatures in the iPSCs, iMGLs, and stimulated iMGLs with IFNγ and demonstrated experimental evidence of epitopes derived from Alzheimer’s and related dementias (ADRD) risk factor proteins being presented on iMGLs. Overall, our characterization of the HLA immunopeptidome within iMGLs provides possible mechanisms behind the progression of ADRD, the unveiling of unique HLA epitope landscapes within iMGLs, and the development of potential immunotherapies.

## Results

### Overview of Study Workflow

In this study, we utilized an iMGLs model derived from the well-characterized KOLF2.1J, the reference line of iPSC Neurodegenerative Disease Initiative (iNDI) project [16, 17]. HLA class I and II proteins and their binding peptides were co-immunoprecipitated. The HLA peptides were eluted under mild conditions, and HLA complexes were characterized subsequently. To investigate how cell type and stimulation alter the immunopeptidome landscape in isogenic cell lines, we included the parental iPSCs, iMGLs, and iMGLs treated with IFNγ. In addition, we characterized the whole-cell proteome, immunopeptidome, and interactome of each cell type using MS-based proteomics. We established a comprehensive workflow to characterize the epitopes and their binding partner proteins in iMGLs, and this workflow can be used for many large-scale HLA immunopeptidome and protein-protein interaction (PPI) studies **(Figure 1)**.

**Figure 1.**
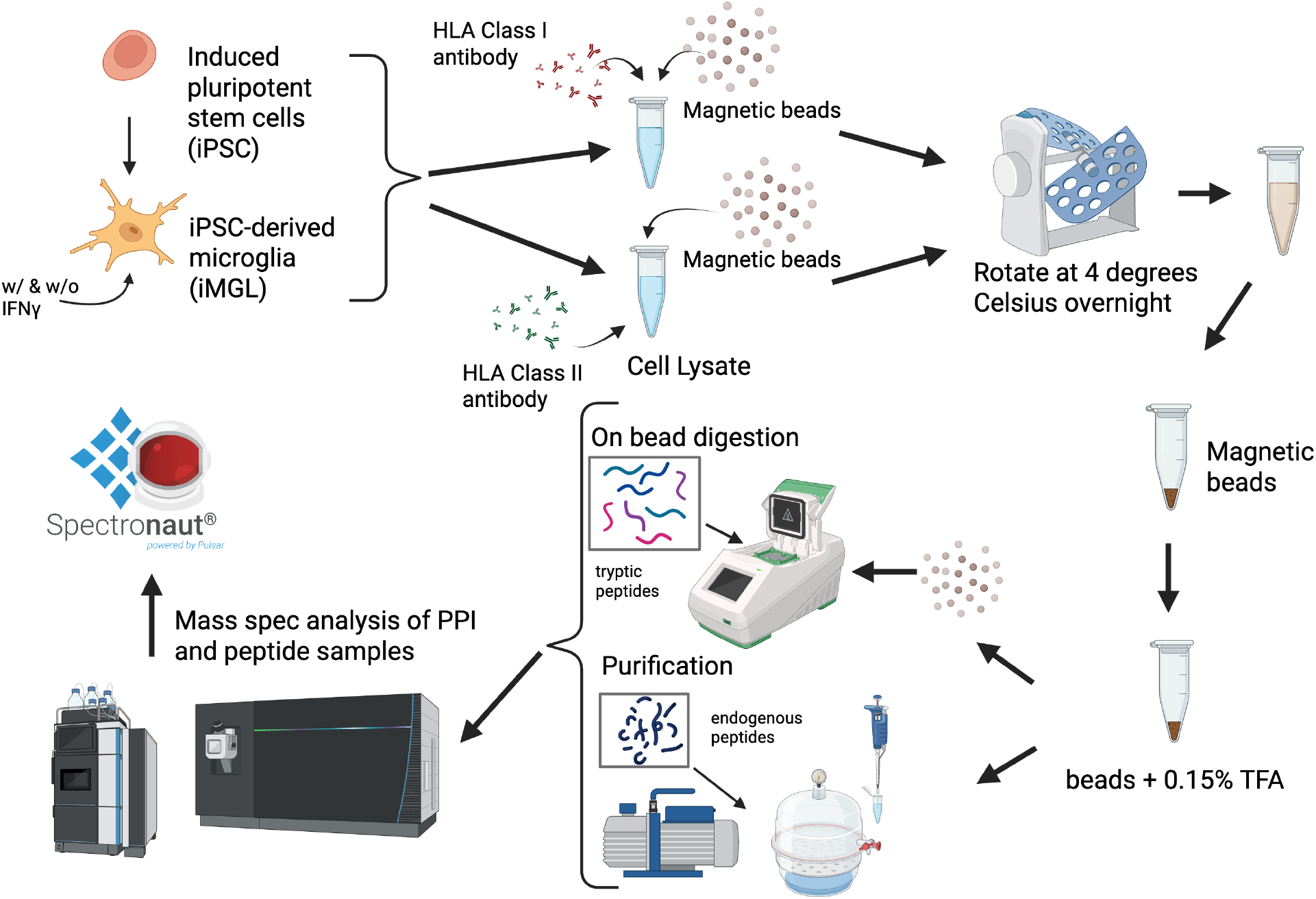
Overview of Study Workflow. The experiment was conducted in KOLF2.1J-based iPSCs and iMGLs which treated with or without IFNγ (n=6). Cells were lysed with a gentle lysing solution. The cell lysate was split into two tubes, and each portion was added with protein A/G magnetic beads and an HLA class I or class II antibody for HLA-I and HLA-II enrichment, respectively. Incubation was conducted overnight at 4 degrees. Cell lysate was removed from the beads and the HLA peptides were eluted in a 0.15% trifluoroacetic acid (TFA) solution for three times. The eluted peptides were collected and freezing dried for DIA-based MS analysis. The separated beads were further processed for HLA interactome profiling using an on-bead digestion with trypsin/lysC followed by LCMS analysis. The MS raw files were database searched using Spectronaut.

### Proteomic characterization of iMGLs

To validate that the iMGL model reflects the characteristics of mature microglia cells, we systematically characterized our iMGLs cells using microscopy and proteomics. Imaging of iPSCs and iMGLs treated with or without IFNγ stained with CD45 and IBA1 demonstrated that these microglial markers were selectively present on iMGLs with or without IFNγ treatment and not on iPSCs (**Figure 2A)**. We performed whole-cell proteomics in iPSCs and iMGLs treated with or without IFNγ and revealed a significant upregulation of various HLA proteins and microglial markers in iMGLs compared to iPSCs (**Table S1**). We used log_2_ fold change (LFC) to quantify effect sizes and an adjusted p-value to determine significance. We observed significant upregulation of HLA molecules (e.g., HLA-I molecules [LFC range = 1.25-2.52, p-value range = 6.10 E-7 to 4.95 E-8] and HLA-II molecules [LFC range = 2.18-16.50, p-value range = 2.47 E-7 to 8.05 E-15] and microglia markers (e.g., PTPRC [also known as CD45, LFC=4.90, p-value = 5.67 E-4], ITGAM [also known as CD11b, LFC=5.93, p-value = 1.20 E-3], CD40 [LFC=3.67, p-value = 5.35 E-9], AIF1 [LFC=14.49, p-value = 1.91 E-11]) in iMGLs compared to iPSCs (**Figure 2B**). In the iPSCs, we observed that HLA-I molecules are expressed significantly higher than HLA-II molecules and both HLA classes are expressed at moderate to high level (**Figure S2**). Furthermore, we performed pathway enrichment analysis of proteins that were expressed at higher levels in iMGLs compared to iPSCs (**Figure 2C**). Notably, within biological process Gene Ontology (GO) terms, we observed proteins involved in immunological responses enriched in iMGLs, such as T-cell activation, antigen processing and presentation, and inflammatory response (p-value range = 1.18 E-19 to 9.6 E-3). Similarly, within cellular component GO terms we observed enriched pathways of upregulated proteins involved in secretory vesicles and MHC protein complexes. When examining the effect of IFNγ treatment in the total proteome of iMGLs, we observed an upregulation of HLA-I molecules (LFC range 0.42-1.94, p-value range = 0.02 to 7.02 E-7) and HLA-II molecules (LFC range 0.47-1.42, p-value range = 7.00 E-4 to 1.16 E-5) in the IFNγ-treated iMGLs **(Figure 2D)**. Importantly, significant upregulation (adjusted p-value <0.001) of HLA-DRA, HLA-DRB1, and HLA-DRB5 were observed upon IFNγ stimulation, indicting the stimulated antigen presentation signaling (**Figure 2E**) [18]. The microglial marker, CD45 displayed significantly increased expression (p-value = 5.19 E-5) upon IFNγ stimulation. These upregulated proteins have been displayed in both mature microglia and activated microglia in the brain. In the pathway enrichment analysis, iMGLs with IFNγ-treatment exhibited an upregulation in proteins involved in immune response, antigen and T-cell receptor signaling, Golgi transport vesicle, HLA protein complex, HLA class I and class II protein binding, and signaling receptor binding (**Figure 2F**). Taken together, our results demonstrated that this iPSC-derived iMGLs model has a similar molecular signature as mature microglia.

**Figure 2.**
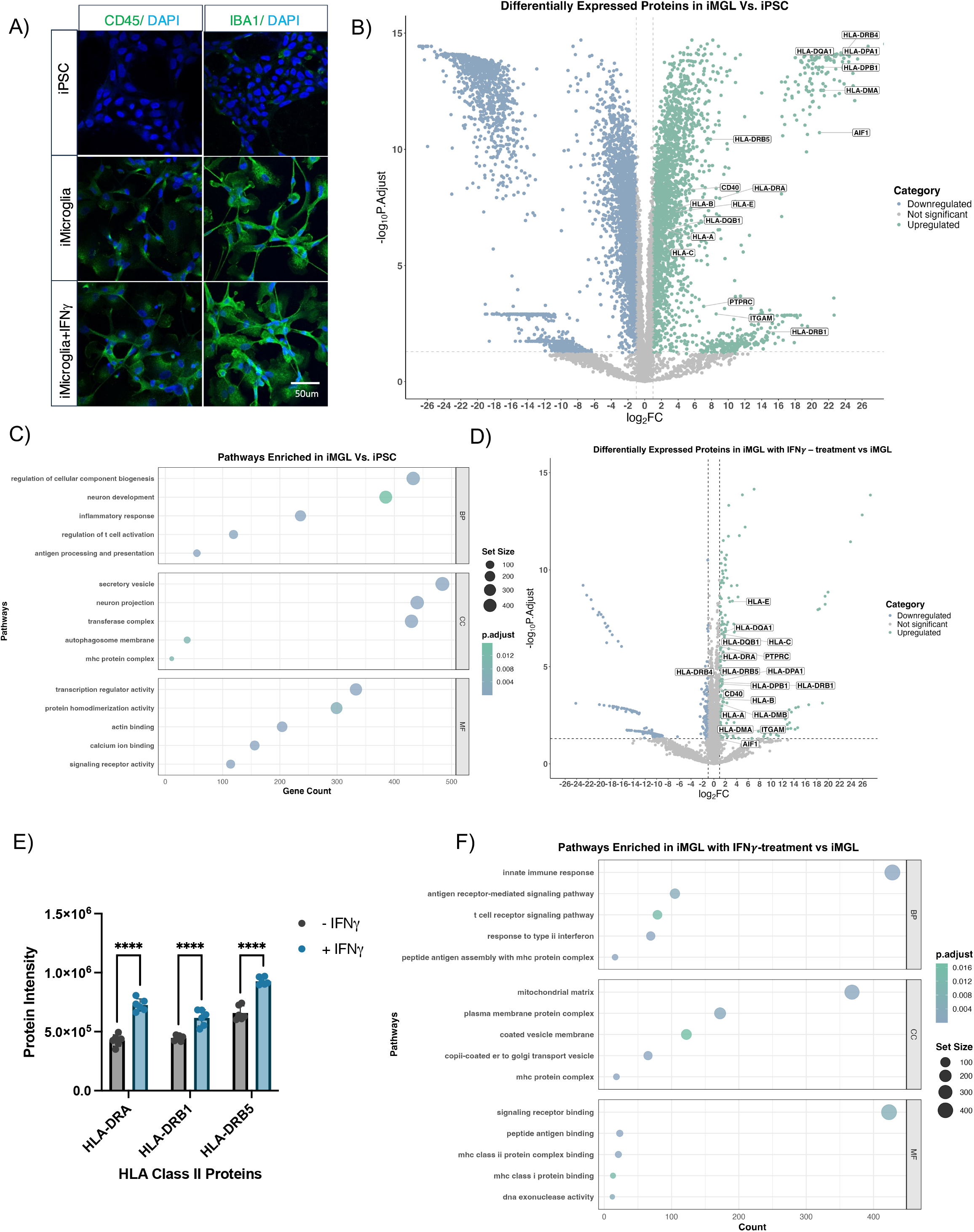
Proteomic characterization of iPSC-derived microglia. **(A)** Immunofluorescence staining of iPSCs, iMGLs, and iMGLs with IFNγ-treatment were stained with the following microglial markers (CD45, IBA1) and nuclear marker (DAPI) displays proper iPSCs differentiation to iMGLs. The scale bar represents 50 μm. **(B)** Differential protein abundance analysis of iPSCs (left) versus iMGLs (right). The following upregulated HLA proteins and microglia markers (HLA-DRB1, ITGAM, PRPRC, HLA-C, HLA-A, HLA-DQB1, HLA-B, HLA-E, CD40, HLA-DRA, and HLA-DRB5) displayed a log_2_(FC) range between 2.5-14. A small cluster of the following upregulated HLA proteins and microglia markers (AIF1, HLA-DMA, HLA-DPB1, HLA -DQA1, HLA-DPA1, and HLA-DRB4) displayed a log_2_(FC) range between 14-17. **(C)** Pathway enrichment analysis of iMGLs versus iPSCs, including biological processes, cellular compartment, and molecular function. **(D)** Differential protein abundance analysis of iMGLs (left) versus iMGLs treated with IFNγ (right). **(E)** IFNγ stimulation significantly upregulates specific HLA-II proteins (HLA-DRA, HLA-DRB1, HLA-DRB5). Each point on a column represents one sample. Statistical analysis was performed using a two-way ANOVA with a Šidák correction test (**** p < 0.0001, *** p < 0.001, ** p < 0.01, * p < 0.05). **(F)** Pathway enrichment analysis in iMGLs treated with IFNγ versus iMGLs; pathways include biological processes, cellular compartment, and molecular function.

### Comprehensive analysis of immunopeptidome in iPSCs and iMGLs

To analyze the HLA immunopeptidome of iMGLs, iMGLs with IFNγ-treatment, and iPSCs samples, we used an in-house informatic pipeline, Protpipe [19]. We observed a significant distinction between the iMGLs immunopeptidome in comparison to the iPSCs immunopeptidome; the HLA-I and HLA-II immunopeptidomes were clustered separately **(Figure 3A)**. In addition, we noticed that the HLA-II immunopeptidome landscape in iMGLs changed upon IFNγ stimulation **(Figure S3A)**. We generated multiple plots focusing on the HLA immunopeptidome deconvolution. The HLA typing of KOLF2.1J showed 6 major HLA alleles of class I and II, and we input our peptidome data and HLA alleles into NetMHCpan to deconvolute the specific HLA epitopes to their respective HLA alleles (**Figure 3B**). For the HLA-I peptidome, we identified over 3000 peptides in iPSCs and identified nearly 5000 peptides in iMGLs and IFNγ-treated iMGLs, consistent with upregulation of HLA molecules in iMGLs **(Figure 3C, Table S2)**. A similar identification pattern was also observed in HLA-II samples with identifying approximately 3500 peptides for iPSCs and 6000 peptides for iMGLs and IFNγ-treated iMGLs (**Figure 3D**). As expected, the peptide length of deconvoluted HLA-I epitopes ranges from 8-14 amino acids, with the most epitopes being 9 mer peptides. We observed a similar HLA-I peptide length distribution in iPSCs, iMGLs, iMGLs treated with IFNγ **(Figure 3E**). In contrast, we found HLA-II peptide length ranges from 11-20 mer, with 15 mer peptides being most frequently found (**Figure 3F**). With the confirmation of optimal HLA-I peptide length, we used an HLA prediction algorithm module within Protpipe to rank the HLA affinity of each epitope to their respective HLA alleles and display the top three highest ranked epitopes **(Figure 3G)**. The bar chart shown in **Figure S3B** shows many epitopes are bound to the specified HLA alleles along with which HLA alleles are more likely to have a binding motif that interacts with epitopes in comparison to other HLA alleles. HLA-A*11:01 had the most favorable binding motif and interacted with the most epitopes in our data set. In contrast, HLA-B*08:01 had the least favorable binding motif and interacted with the least epitopes. For HLA-II immunopeptidome deconvolution, HLA-DRB1*01:01 and HLA-DPB1*14:01 presented the most and least epitopes, respectively (**Figure S3C**). In summary, our experiment enriched a large number of HLA-I and HLA-II epitopes which display distinct signatures in iPSC and iMGLs, and that more HLA-II peptides were enriched than HLA-I in iMGLs.

**Figure 3.**
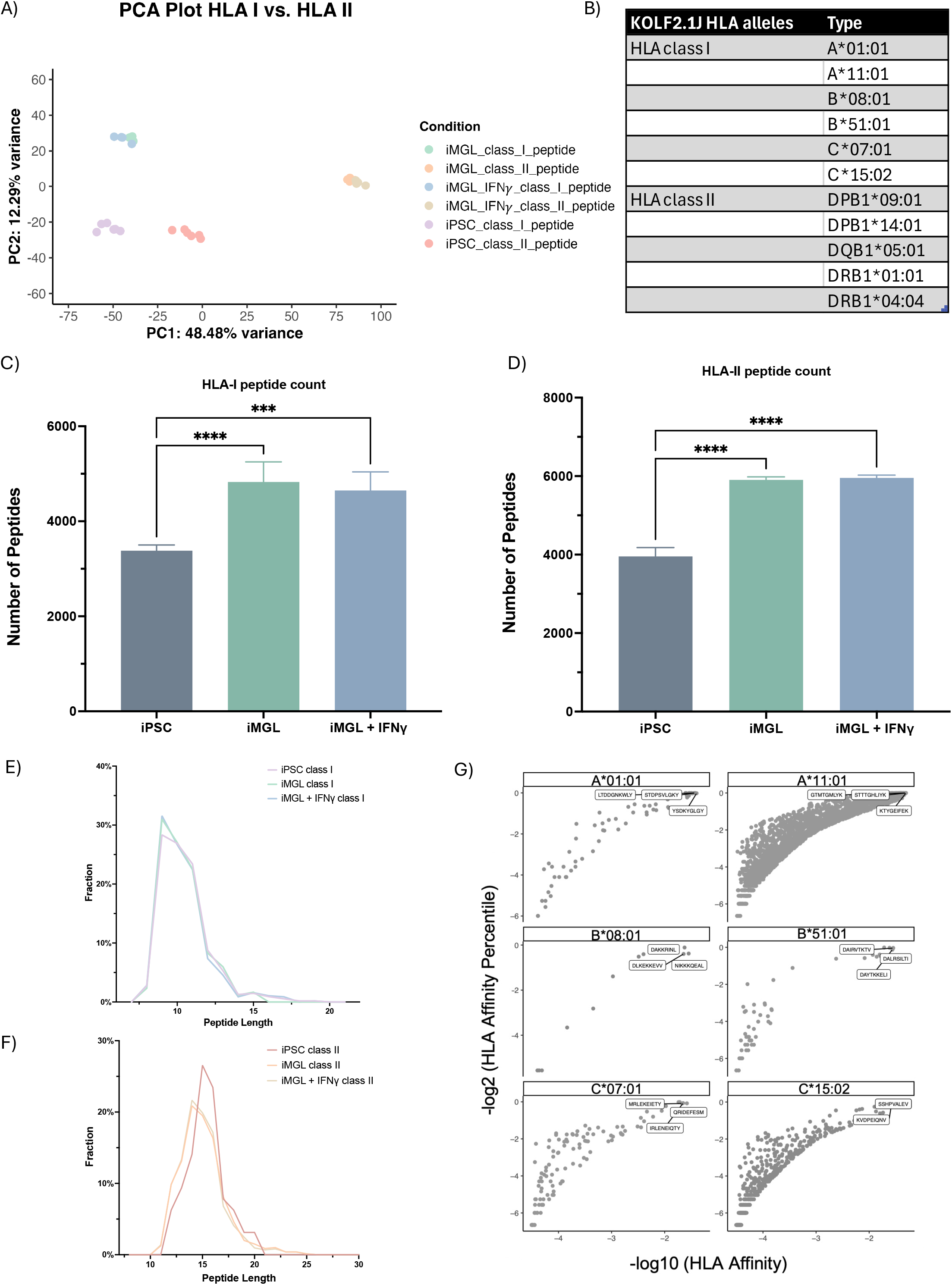
Immunopeptidome characterization of HLA-I and II in iPSCs and iMGLs. **(A)** PCA plot displays the distinction between HLA class I and class II immunopeptidome and a distinction between iPSCs and iMGLs cell types. PC1 presented with 48.48% variance and PC2 presented with 12.29% variance. **(B)** HLA alleles in the KOLF2.1J cell line. This cell line was used for both the iPSCs and iMGLs cells. **(C)** Total peptides count for HLA-I samples in iPSCs, iMGLs, and iMGLs with IFNγ-treatment. Statistical analysis was performed using a one-way ANOVA and Turkey’s multiple comparisons test (**** p < 0.0001, *** p < 0.001, ** p < 0.01, * p < 0.05). Error bars represent standard deviation. **(D)** Total peptides count for HLA-II samples in iPSCs, iMGLs, and iMGLs with IFNγ-treatment. Statistical analysis was performed using a one-way ANOVA and Turkey’s multiple comparisons test (**** p < 0.0001, *** p < 0.001, ** p < 0.01, * p < 0.05). Error bars represent standard deviation. **(E)** Peptide length distribution in HLA-I comparing total peptide count in iPSCs, iMGLs, and iMGLs with IFNγ-treatment. Peptide length was filtered by binding affinity with HLA-I allele, with affinity rank < 2%. **(F)** Peptide length distribution in HLA-II comparing total peptide count in iPSCs, iMGLs, and iMGLs with IFNγ-treatment. Peptide length was filtered by binding affinity with HLA-I allele, with affinity rank< 5%. **(G)** HLA-I deconvolution presents top three binding epitopes for each HLA-I allele with binding affinity threshold cutoff at < 2%.

### HLA epitopes associated to Alzheimer’s disease and related dementias

We sought to discover HLA epitopes derived from ADRD associated genes that we reported previously (**Table S3**) [20]. To assess the specific binding affinity for these epitopes and whether they would interact with the specified HLA alleles from the KOLF2.1J cell line, we applied an HLA prediction algorithm. In iMGLs, we observed 25 epitope sequences that bound within the top 7% to the selected HLA alleles relative to the other epitopes in the data, eight of these sequences presented within the 2% rank criteria. All these 25 epitope sequences were derived from 15 ADRD-risk proteins, such as MAPT (also known as Tau) and amyloid-beta precursor protein (APP) **(Table 1)**. Next, by identifying the presence of epitopes derived from ADRD-associated genes in the HLA immunopeptidome data, we evaluated how changing the epitope sequences to have it contain a pathologic single nucleotide polymorphism (SNP) associated with ADRD-risk would affect the HLA binding affinity. Using NetMHCpan as a prediction algorithm, we selected five ADRD-associated risk proteins (i.e., APP, MAPT, TARDBP, FUS, SNCA) that have pathologic SNPs that lead to the development of AD, frontotemporal lobe dementia (FTLD), amyotrophic lateral sclerosis (ALS), Parkinson’s disease, or Lewy body dementia (LBD). We compared the HLA binding rank of the mutated epitope sequences to the wildtype sequences. We kept the mutated epitope sequences that displayed an equal or greater binding rank compared to their wildtype counterparts to the selected HLA alleles. From that comparison, we predicted 31 mutant epitope sequences that contain an ADRD-risk singular mutation from the five selected ADRD risk genes that exhibited a stronger HLA binding affinity than their wildtype counterpart **(Figure 4)**. These results suggest that a mutated epitope derived from ADRD risk proteins can be presented on the surface of iMGLs and potentially exert immunogenicity.

**Table 1.**
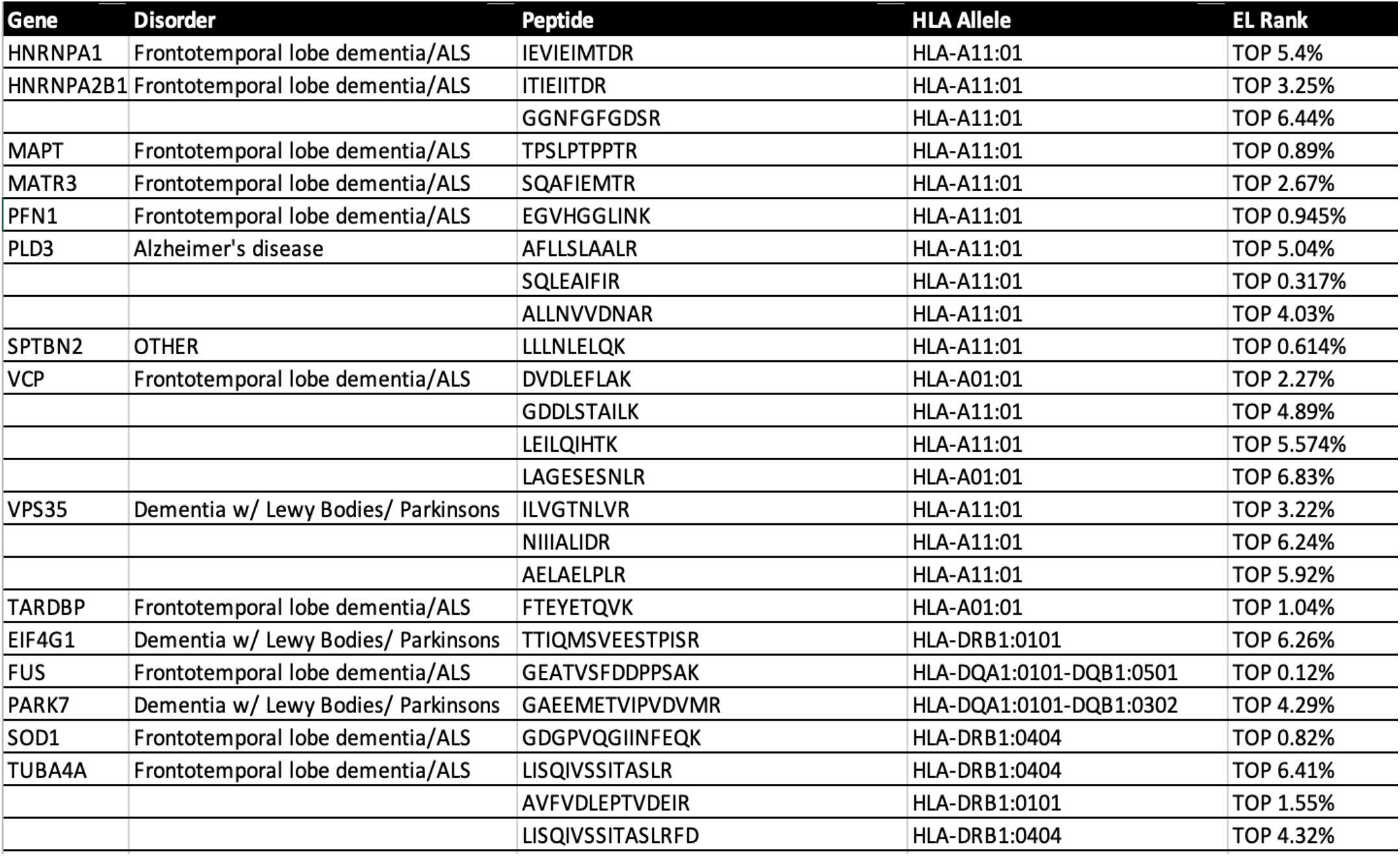
Identification of ADRD risk protein derived HLA epitopes in iMGLs.

**Figure 4.**
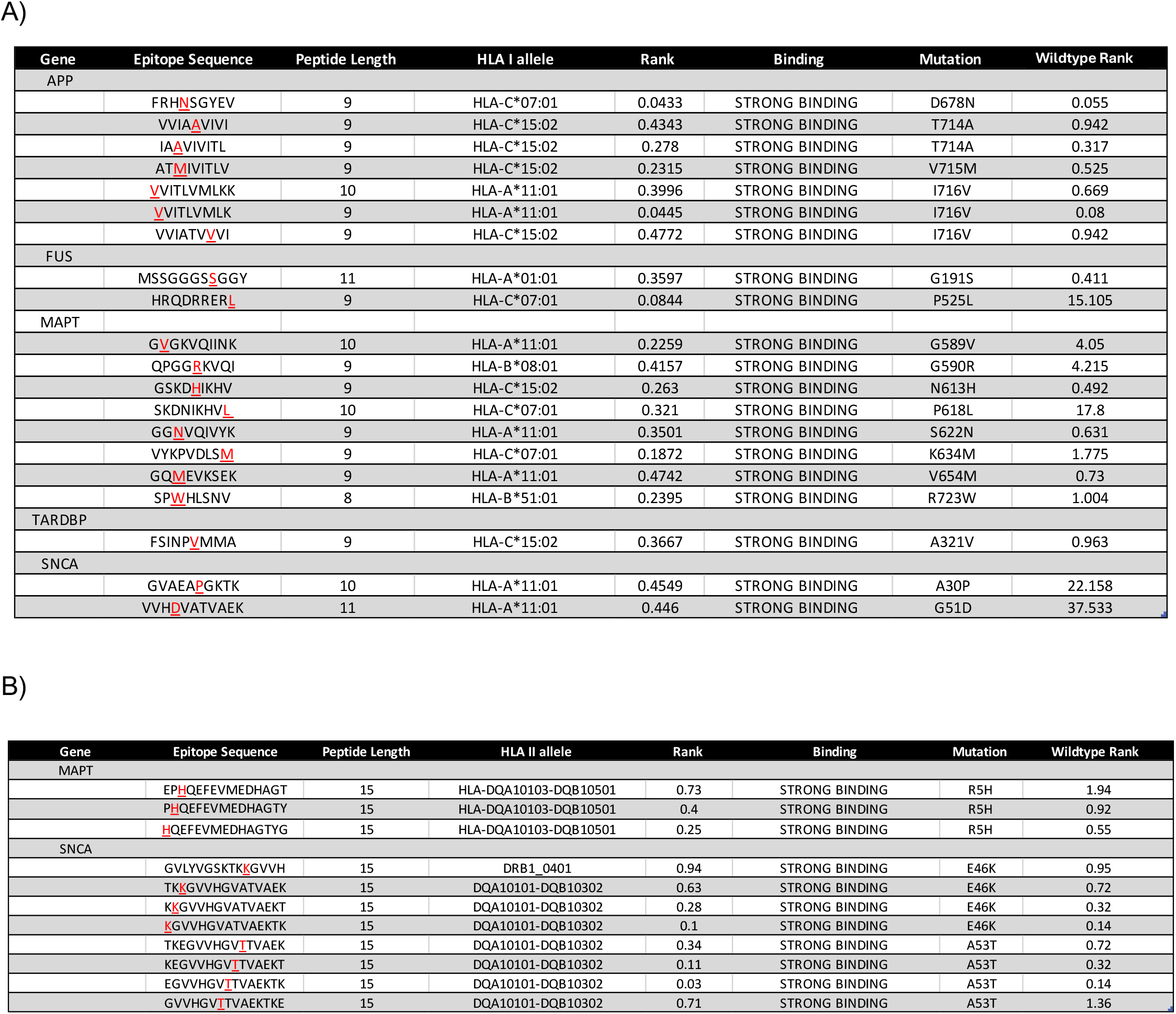
Mutant epitope sequences that contain pathologic ADRD mutation in five common ADRD-associated genes: MAPT, FUS, APP, SNCA, and TARDBP. **(A)** Mutant epitope sequences that strongly interacted with an HLA class I allele from the five common ADRD-associated genes: MAPT, FUS, APP, SNCA, and TARDBP (also known as TDP-43) as strong binder (affinity rank< 0.5%). **(B)** Mutant epitope sequences that strongly interacted with an HLA class II allele from the five common ADRD-associated genes: MAPT, FUS, APP, SNCA, and TARDBP as strong binders (affinity rank< 1%)

### Profiling of HLA interactome in iPSC and iMGLs

Following immunopeptidome analysis, we next evaluated the PPIs associated with HLA. For this, we performed an on-bead digestion with the magnetic beads used in the HLA co-immunoprecipitation followed by MS-based proteomics analysis. In the HLA PPI complex, we observed that iMGLs and iPSCs share 37% of interacting proteins, 7% of proteins are unique to iMGLs, and 55% were unique to iPSCs (**Figure 5A)**. Unexpectedly, with iPSCs displaying more interaction proteins, we analyzed the unique interaction proteins in each cell line. In iPSCs, the GO biological processes of the unique proteins revealed processes involved in metabolic processes, cellular biogenesis, and cytoskeleton organization **(Figure S4A)**. In contrast, the unique interaction proteins to iMGLs revealed processes involved in regulation of immune response, phagocytosis, and leukocyte activation **(Figure S4B)**. With distinguishing the interaction protein unique to cell type, we evaluated how HLA class affected interaction proteins. In both iPSCs and iMGLs, we observed a large amount of shared interaction proteins in HLA-I and HLA-II, with minimal unique interaction proteins to class type **(Figure 5B)**. Similarly, we observed immune responsive proteins are recruited in the HLA complex in iMGLs based on GO pathway analysis of unique PPIs of HLA-I or HLA-II in iMGLs (**Figure S4C and S4D**) and in iPSCs (**Figure S4E ana S4F**), respectively. When comparing classes and cells type, many proteins showed elevated interaction with HLA-I and HLA-II in iMGLs compared to iPSCs; for example, TGM2 (LFC = 17.42, p-value = 2.54 E-10), SQOR (LFC = 18.06, p-value = 5.66 E-10), FERMT3 (LFC = 16.71, p-value = 1.55 E-9), PLEK (LFC = 15.55, p-value = 1.13 E-8), and B2M (LFC = 16.15, p-value = 1.77 E-8) with HLA-I (**Figure 5C**) and REP15 (LFC = 16.14, p-value = 1.47 E-8), B2M (LFC = 16.73, p-value = 2.13 E-8), IFI16 (LFC = 16.19, p-value = 2.10 E-9), and LAMB3 (LFC = 18.12, p-value = 1.21 E-7) with HLA-II (**Figure 5D**). GO enrichment analysis of HLA-I complex demonstrated increased activation of pathways involved in phagocytic and endocytic vesicle membranes, and humoral immune response in iMGLs while displaying a suppression in RNA processing and splicing, and mRNA processing (p-value range = 6.74 E-8 to 1.4 E-4) (**Figure S5A**). In addition, the KEGG database analysis revealed interacting proteins involved in increased activation of antigen processing and presentation and lysosomal activation (p-value range = 7.52 E-6 to 4.30 E-2) (**Figure S5B**). For HLA-II complex, GO analysis revealed increased activation of endocytosis, vesicle membrane, and lysosome in iMGLs (p-value = 5.44 E-9) (**Figure S5C**) and these processes also appeared to be activated in the KEGG database (p-value range = 1.00 E-10 to 3.10 E-3) (**Figure S5D**). We further visualized enriched HLA interaction partners enriched in iPSCs and iMGLs treated with or without IFNγ, showing eleven unique HLA-II interaction proteins significantly enriched in most of iMGLs HLA-II samples but not in the HLA-I iMGLs or iPSCs (i.e., ACACB, WAS, ADPRH (LFC = 9.86, p-value = 0.016), PDAP1(LFC = 12.15, p-value = 9.00 E-3), ACAP2, PDSS1 (LFC = 12.37, p-value = 9.00 E-3), CCDC117 (p-value = 9.00 E-3), APIP (p-value = 9.00 E-3), PRDM6 (LFC=15.04, p-value= 9.00 E-3) (**Figure 5E)**. Our results highlight the unique HLA binding protein in different cell types (i.e., iPSCs versus iMGLs) and microglial states.

**Figure 5.**
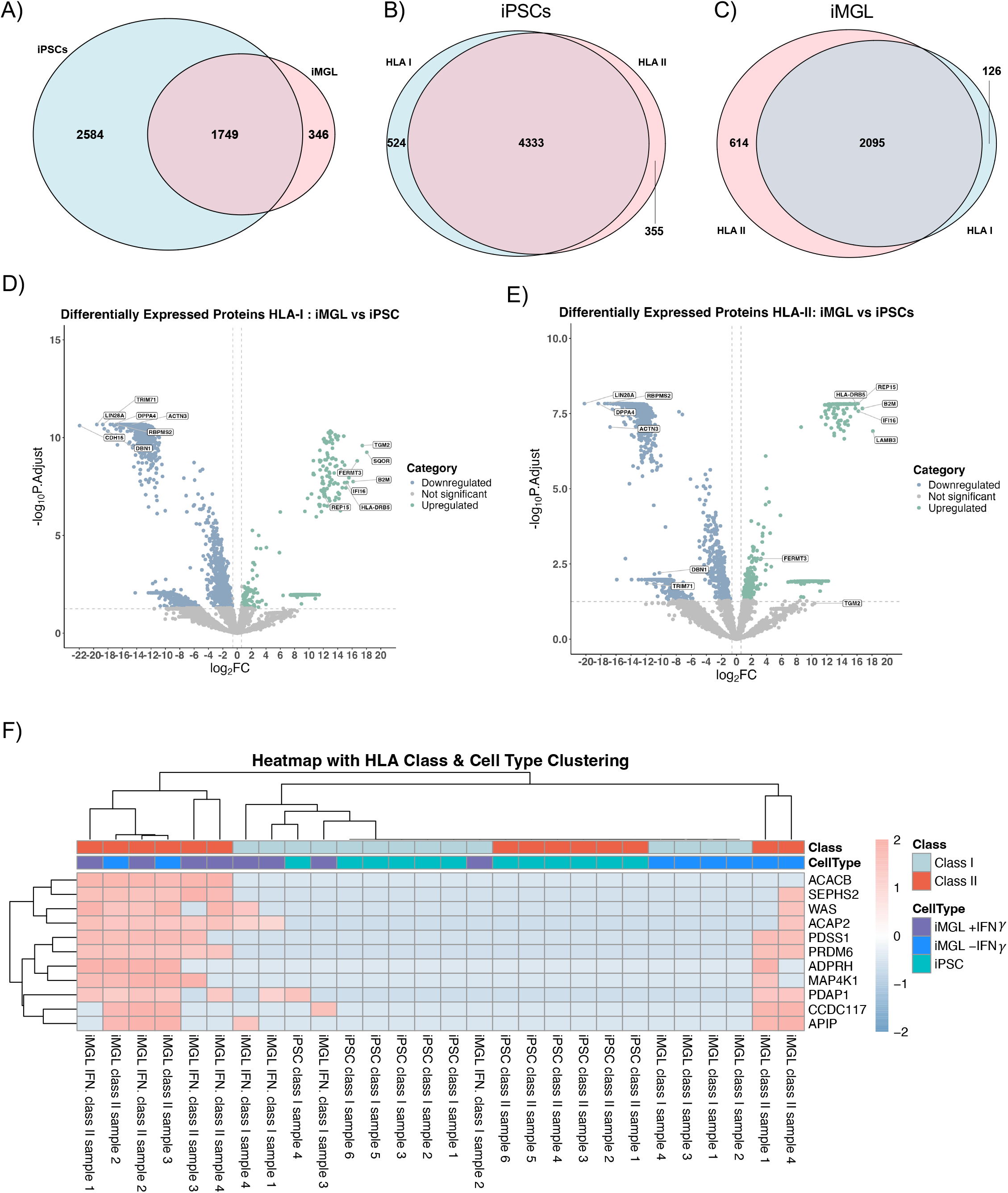
Distinction between interaction proteins in iPSCs vs iMGLs exhibited proteins and pathways unique to each cell type and HLA class. **(A)** Venn diagram showing iPSCs and iMGLs share 1749 HLA interacting proteins of which 346 interacting proteins were unique to iMGLs whereas 2584 interacting proteins were unique to iPSCs. **(B)** Left Venn diagram showing iPSCs and iMGLs share 4333 HLA-I interacting proteins, and right Venn diagram showing iPSCs and iMGLs share 2095 HLA-II interacting proteins. **(C)** Differential abundance analysis of HLA-I interacting proteins in iMGLs compared to iPSCs. The following upregulated proteins (TGM2, SQOR, FERMT3, PLEK, and B2M) displayed a log_2_(FC) range between 11-16. The significantly downregulated proteins (DBN1, RBPMS2, ACTN3, DPPA4, TRIM71, LIN28A, and CDH15) displayed a log_2_(FC) range between 15-22. **(D)** Differential abundance analysis of HLA-II interacting proteins in iMGLs compared to iPSCs. The following HLA-II interacting proteins (HLA-DRB5, REP15, B2M, IFI16, and LAMB3) were upregulated with a log_2_(FC) range between 17-19. The following downregulated proteins (DBN1, TRIM71, RBPMS2, LIN28A, DPPA4, ACTN3) exhibited a log_2_(FC) range between 11-20. **(E)** Heat map displaying top enriched HLA II interacting proteins in iMGLs. Heatmap was scaled using row z-scores.

## Discussion

Inspired by the critical role of immunopeptides in T-cell activation and immunotherapy, we sought to provide the very first HLA peptidome profiling in microglia, the major antigen-presenting cells of the CNS. This rich resource is useful to deepen our understanding in brain immunology and support the possibility that HLA epitopes may play a role in neurodegenerative disease, and if so, could lead to possible therapeutic interventions. These findings highlight the unique immunopeptidome landscape in iMGLs and its potential to present peptides derived from ADRD risk genes and possible links to neurodegeneration.

As expected, the total proteome of iMGLs versus iPSCs were highly distinct. iMGLs expresses multiple HLA class II proteins not found in iPSCs, along with overall upregulation in HLA class I and II proteins, consistent with prior studies [8, 21]. Along with significant upregulation of HLA proteins, we detected other common microglia makers (e.g., CD45), indicating appropriate differentiation of iPSCs to iMGLs, consistent with the markers recognized in the human microglia proteomic landscape [22]. Furthermore, we observed enrichment in HLA proteins with IFNγ stimulation and the positive enhancement of immune pathways such as innate immune response, HLA antigen processing and presentation, and T-cell signaling, similar to prior reports [14, 23].

Through using a systematic HLA peptide enrichment workflow, high-resolution MS, and HLA binding prediction algorithms, we provide the most comprehensive HLA peptidome in microglia cells to date. Our immunopeptidome analysis found different landscapes of iPSCs, iMGLs with and without IFNγ stimulation, suggesting that antigen presentations are altered by cell type and state. The variation of the immunopeptidome between HLA classes and cell type indicates variation in peptide sequences found interacting with HLA in iMGLs compared to iPSCs. In addition, we noticed variation between HLA classes, agreeing with the specificity of the HLA-I and HLA-II binding grooves. To properly confirm interaction of HLA-peptide complexes, we examined the peptide length in our data. We observed that the majority of our peptides for HLA-I were 9 mer and for HLA-II were 14-15 mer, which aligns with the binding groove characteristics of each class [7]. Analyzing the eluted epitope sequences, we noted the identification of epitopes derived from ADRD-risk factor proteins along with proteins involved in immunological and inflammatory signaling (e.g., PSMD2, EEF1A1, PTMA, TLN1) [24-27]. Leveraging the HLA prediction algorithm to explore the possibility that ADRD risk variants containing epitope may have the opportunity to be presented, we discovered 31 epitopes containing common pathogenic variants from SNCA, APP, MAPT, TDP-43, and FUS, indicating that mutated epitope sequences in ADRD associated risk genes can potentially bind and be presented by HLA. These results from our immunopeptidome analysis are consistent with findings investigating the antigen presenting mechanisms within microglia [1, 28].

We observed very different HLA interactome landscape in iMGLs compared to iPSCs; mainly, interaction proteins involved in immunological responses and antigen processing and presentation are significantly enriched in iMGLs. Our findings showed the presence of few proteins specific to either HLA-I or HLA-II, suggesting a great overlapping interactome of both HLA-I and HLA-II. Specifically, the proteins B2M, HLA-DRB5, IFI16, FERMT3, and REP15 were enriched in both classes and display involvement in assertions of CD8+T cells responses, chemokine production, receptors recycling, interferon production, and antigen presentation [29-34]. PPI network analysis (**Figure S6**) displays various interactions between both HLA-I and HLA-II proteins supporting findings of HLA proteins cross-linking and interacting [35]. With these similarities in protein interactions, we noticed related biological processes between these two classes, although they present with differing antigen processing mechanisms. Furthermore, HLA-II exhibited eleven significant interaction proteins, of which, WAS, ADPRH and ACAP2 were specifically involved in vesicle trafficking, modulation of inflammatory pathways, and regulation of cytoskeleton [36-38]. The HLA-II processing mechanism uptakes extracellular proteins via phagocytosis and which becomes degraded in endosomes and lysosomes [39]. Peptides are sorted in vesicles where they are then brought to HLA-II proteins. The pathways of these unique interaction proteins to HLA-II in iMGLs display high involvement in the activation of endocytosis and vesicular trafficking along with fc gamma R-mediated phagocytosis. In the IFNγ induced iMGLs, we observed HLA-II had a significant enrichment of interacting proteins involved in immune response and antigen pathways.

Therefore, our results suggested a possible HLA centered mechanism in microglia which plays a key role in CNS adaptive immune response and possible link to neurodegeneration. This comprehensive profiling of the HLA immunopeptidome and interactome of iMGLs explored the potential mechanisms driving the progression of ADRD, revealing potential biomarkers and highlighting the significant partnership of HLA-II in a pathological state. Understanding the critical role of HLA epitope landscape in microglia provides the investigation and development of potential immunotherapies.

## Methods

### Tissue Culture

#### iPSCs culture

The cell line for iPSCs was KOLF2.1J (Jackson Laboratory Cat. #JIPSC1000). iPSCs were thawed into a single well of a Matrigel (1:100, Corning Cat. #354277) coated 6-well plate in Essential 8™ medium (Thermo Fisher Cat. #A1517001) with Chroman-1 (1:1000, MedChemExpress Cat. #HY-15392). After 24 hours, the medium was changed to Essential 8™ medium. For passaging, cells were dissociated with 0.5 mM EDTA-PBS solution (EDTA; 1:1000, Thermo Fischer Cat. #AM9260G) (Phosphate Buffered Saline; Lonza Cat. #17-516F). Dissociated iPSCs were plated onto Matrigel-coated 10 cm plate (Corning Cat. #430167) at 700,000 cells/plate in Essential 8™ medium with Chroman-1. After 24 hours, the medium was changed to Essential 8™ medium. Cells were kept in an incubator at 37 °C with daily Essential 8™ medium changes.

#### iPSC-derived microglia differentiation

iPSCs were cultured in Essential 8™ medium without ROCK inhibitor for at least 2 days prior to initiating differentiation. On the first day of differentiation, 10,000 cells were plated per well in a V-bottom 96-well ultra-low attachment plate (Sanko) in Embryoid Body Medium (EBM) containing Essential 8™ medium, 10 μM ROCK inhibitor, 50 ng/mL BMP-4, 20 ng/mL SCF, and 50 ng/mL VEGF-121. The following day, 80 μL of medium was removed from each well and replaced with 100 μL of EBM without ROCK inhibitor. EBs were cultured for an additional 2 days with daily medium changes using fresh EBM without ROCK inhibitor.

On Day 4, the EBs were transferred from the 96-well plate to a 50 mL tube using wide-orifice pipette tips (P200) or 1 mL precut tips. EBs are generally visible and will settle at the bottom of the tube. Most of the EBM medium was removed, and approximately 200 EBs were transferred to a T75 flask coated overnight with 10 μg/mL Fibronectin. To each flask, 20 mL of Microglia Progenitor Media (MPM) was added, consisting of X-VIVO™ 15 (Lonza), 2 mM GlutaMax™, 55 μM β-mercaptoethanol, 100 ng/mL M-CSF, and 25 ng/mL IL-3. EBs were allowed to adhere over the next 3-4 days, with media changes every other day (⅔ media change each time). If the EBs failed to adhere, differentiation continued, as the floating EBs still produced microglia progenitor cells.

Large amounts of microglia progenitor cells typically appeared between Day 12 and Day 18. The microglia progenitor cells were collected by centrifugation, and approximately 4 million cells per dish were seeded into non-coated 10 cm plates for the final differentiation in Microglia Maturation Media (MMM), which consisted of Advanced RPMI, 2 mM GlutaMax™, 100 ng/mL IL-34, and 10 ng/mL GM-CSF. The cells were further differentiated for an additional 10 days, with full MMM media changes every other day. On Day 10, microglia cells were treated with 25 ng/mL IFNγ for 24 hours.

### Immunocytochemistry

iPSCs and iMGLs cells were grown on Poly-D-Lysine/Laminin coated 12mm glass coverslip (354087, Fisher Scientific) overnight. For the staining of microglia-specific markers, cells were fixed with 4% PFA for 12 minutes, washed in PBS at room temperature, and incubated for 1 hour in blocking buffer (5% Normal Donkey Serum (NDS), 0.1% Triton X-100 in PBS). Afterwards, primary antibodies (rabbit anti-CD45 and IBA1) were used at 1:250 and incubated overnight in 1% NDS, and 0.1% Triton X-100 in PBS at 4°C. The next day, cells were washed three times for 10 minutes with PBS and incubated with 1:500 Alexa Fluor conjugated secondary antibody (anti-goat rabbit 488; Invitrogen) for 1 hour at RT. After three washes with PBS, cells were mounted on glass slides (Fisher Scientific) coverslips using Prolong Gold Antifade mounting media (P10144, Invitrogen), and imaged using a Zeiss LSM 880 confocal microscope equipped with Plan-Apochromat 63X/1.4 numerical aperture oil-objective (Carl Zeiss AG).

### HLA Immunoprecipitation

iMGL or iPSCs in 10 cm plates with ∼ 90-95% confluency was washed three times with ice cold PBS. Cells were lysed with 2 mL of lysis buffer (0.05M Tris-HCl [pH 8.0] [Thermo Scientific Cat. #15575020], 0.15 M NaCl [Thermo Scientific Cat. #24740011], 1% TritonX-100 [MilliporeSigma Cat. #X100], LC/MS grade water [Thermo Scientific Cat. #51140], and 1x EASYpack Protease Inhibitor Tablet [MilliporeSigma Cat. #5892970001]). Using a cell scraper to lift the cells, 1.5 mL of cell lysate was transferred into an Eppendorf tube and then rotated for 30 minutes at 4 °C. Lysates were then centrifuged at 18,213 rcf for 30 minutes at 4 °C. Leaving the cell pellet undisturbed, supernatants were transferred to new Eppendorf tubes.

Immunoprecipitation for HLA-I or HLA-II were performed as follows. For each sample, 50 μL of Protein A/G magnetic beads (ThermoFisher Cat. #88803) and 25 μl of HLA antibody (HLA-I antibody: BioXcell Cat. #BE0079; HLA-II antibody: Cell Signaling Technology Cat. #68258S) are used. The Protein A/G magnetic beads were prepped by being washed three times with an IP wash buffer (500mL of DPBS [ThermoFisher Scientific Cat. #14190144] with 0.01% TritonX-100 [MilliporeSigma Cat. #X100]). Beads were resuspended in 50uL of IP wash buffer per each sample. 25uL of HLA antibody was directly added to the samples after the addition of the beads. Lysates were rotated overnight at 4 °C. After incubation, samples were placed on a magnetic stand, the supernatant was collected and stored in -80 °C. The magnetic beads were washed three times with 1 mL of IP buffer wash and three times with 1 mL of DPBS (ThermoFisher Scientific Cat. #14190144). The beads were then washed once with LC/MS grade water (Thermo Scientific Cat. #51140).

### Peptidome enrichment and purification

For HLA peptides elution, beads were soaked three times in 200 μL of 0.15% TFA for 5 minutes. Supernatant was placed in a new Eppendorf tube. For purification, the supernatant was transferred to tC18 50 mg Sep-Pak cartridges (Waters Cat. #WAT200685) attached to a Hypersep Glass Vacuum Manifold (ThermoFisher Cat. #60104-233). To prep the cartridges, 200 uL of 100% methanol (Millipore Sigma Cat. #1060351000) was added twice, 100uL of 99% ACN /0.1% FA was added once, and 500uL of 1% FA (ThermoFischer Scientific Cat. #28905) was added four times. Solvents that were eluted were collected in a 15mL conical tube and discarded after prep.

Peptides were eluted via cartridge using 250 μL of 15% ACN (Thermo Scientific Cat. #PI51101) /1% FA (ThermoFischer Scientific Cat. #28905) and twice with 250 μL of 50% ACN/ 1% FA and collected in a new Eppendorf tube. Peptides were then dried and stored at -80 °C.

### Protein-Protein Interaction

A trypsin digest was performed on the remaining magnetics beads left in the original Eppendorf tubes that were used in the pre-elution. For this digest, 1 μg of Trypsin/Lys-C (Promega Cat. #V5073) and 100 μL of 50 mM ammonium bicarbonate (MilliporeSigma Cat. #A6141) was needed per sample. 100uL of digest was added to each sample. These samples were placed in a thermocycler and incubated at 37 °C for 16 hours. After incubation, samples were placed on a magnetic stand and the supernatant was collected in new Eppendorf tubes. Samples were dried and stored at -80 °C.

### Liquid Chromatography Mass Spectrometry Analysis

Dried peptides were reconstituted using 2% acetonitrile (ACN) (Thermo Scientific Cat. #PI51101) in 0.4% trifluoroacetic acid (TFA) (Thermo Scientific Cat. #85183) and normalized to 0.2 μg/μL then centrifuged for 15 minutes at 4 °C. The liquid chromatograph/mass spectrometry system (LC/MS) we used for analysis was a hybrid Orbitrap Eclipse mass spectrometer and UltiMate 3000 nano-HPLC system. Liquid phases A (5% DMSO [Sigma-Aldrich Cat. #D2650] in 0.1% formic acid (FA) in water) and B (5% DMSO in 0.1% FA in ACN) were used. Peptides were separated on a ES903 nano column (75 μM x 500 mm, 2 μM C18 particle) using a 2-hour linear gradient. Flow rate was set constant at 300 nL/min (0-5 min [2% liquid phase B], 5-120 min [35%-80% liquid phase B], 120-125 min [35%-80% liquid phase B], 125-135 min [80% liquid phase B], 135-136 min [80%-2% liquid phase B], 136-150 min [2% liquid phase B]. Samples were then loaded on a trap column with the loading pump set at a constant rate of 5 μL/min. For fragmentation, high collision dissociation with 30% collision energy was used. We performed a data-independent acquisition (DIA) method with no FAIMS. The MS1 resolution was set to 120k and MS2 resolution was set to 30K. Precursor range was set to 400-1000 m/z with the isolation window at 8 m/z resulting in 75 windows per each scan cycle for MS2 scanning. The scan range for MS2 was specified as 145-1450 m/z with the loop control being set to 3 seconds.

### Database Search

Mass spectra results were input into Spectronaut software, version 19.4 (Biognosys) using the UniProt human proteome reference FASTA file (UP000005640) which contains 20,420 protein entries. HLA classes were separated at search.

#### HLA Peptidome Database Search

Peptide files were performed as a non-specific search and no digestion enzyme was selected. No fixed modifications were selected though for variable modifications, N-terminal acetylation and oxidation were selected. For HLA-I, the peptide length was set between 7 to 20 amino acid lengths. For HLA-II, the peptide length was set between 7-30 amino acid lengths. False discovery rate (FDR) for PSM, peptide, and protein groups was set to 0.01. We did not perform cross-run normalization.

#### HLA Protein-Protein Interaction Database Search

Protein files were performed as a specific search with Trypsin/P and LysC/P selected for digestive enzymes. Carbamidomethylation was selected as a fixed modification. N-terminal acetylation and oxidation were selected as variable modifications. FDR for PSM, peptide, and protein groups was set as 0.01. Cross-run normalization was not performed.

### Bioinformatic analysis

We used the functional pipeline Protpipe for data analysis. Differential intensity was conducted performing limma with the log2 fold change being set at a minimum of 2 and the p-value adjusted to proper threshold to determine the significantly different proteins. Enrichment pathway analysis applied the Gene Ontology database to define specific proteins based on their biological process (BP), cellular compartment (CC), and molecular function (MF). Using the GraphPad Prism software (Version 10.3), we applied two-way analysis of variance (ANOVA) analyses. The PCA, pathway enrichment, upset plot, heatmap, and bar charts were performed using the ggplot2 package from Rstudio (Version 4.2).

#### HLA typing

HLA typing was determined by HLA genotyping from the whole genome sequencing of KOLF2.1J iPSC line (https://github.com/cory-weller/KOLF2.1-HLA).

#### HLA peptide prediction

For HLA peptide affinity prediction that were used for peptide distribution and ADRD-risk sequence predictions we applied the prediction algorithm NetMHCpan (version 4.1 for HLA-I and version 4.3 for HLA-II). The interactions are determined by the specific binding motif for each HLA allele and their specified essential anchor residues. NetMHCpan is an artificial neural network (ANN) and generates peptide-binding predictions using motif search strategies or statistical matrices determine the frequency of each amino acid in each position [40]. The input peptide sequences were presented in the ANN in three manners: conventional sparse encoding, Blosum encoding, and a mix of both methods. Models were trained with stochastic gradient descent using back propagation [41]. For statistical testing for the ANN model, they performed cross-validation with one tailed binomial test and standard significance at 0.05. For stringent filtering, we set affinity rank score cutoff at < 2% for HLA-I and < 5% cutoff for HLA-II. Strong binders for HLA alleles were cut off at the affinity rank of < 0.5% for HLA-I and < 1% for HLA-II. For ADRD-risk mutations, pathological single point mutations were selected from the UniProt variant viewer source. FASTA file of ADRD risk-gene protein sequence was inputted into NetMHCpan to get rank of the non-pathologic sequences. Pathologic single point ADRD mutations were then inputted to the FASTA file, and the new mutated sequence HLA affinity rank was compared to the normal sequence rank. Mutated sequence was kept if it presented with a lower rank compared to the non-mutated sequence. HLA peptide deconvolution analysis was conducted in Protipipe to determine potential epitope sequences and their binding affinity with an HLA-I allele. For proper filtering, the affinity score cutoff was set at < 2%.

## Data Availability

All raw files from mass spectrometry analysis have been deposited to the ProteomeXchange Consortium through the PRIDE partner repository with the dataset identifier PXD061309.

## Acknowledgements

This research was supported in part by the Intramural Research Program of the NIH, National Institute on Aging (NIA), National Institutes of Health, Department of Health and Human Services; Project number ZIAAG000536.

## Author Contributions

Conceptualization, S.K., A.B., J.H.P., Y.A.Q.

Methodology, S.K., A.B., J.H.P., Y.A.Q.;

Data curation, Z.L., C.T., C.A.W;

Formal analysis, Z.L., C.A.W., C.T.;

Investigation, S.K., A.B., Y.H., J.H.P., Z.L., M.I., C.T., B.J., J.E., I.K., N.C.,Y.A.Q.;

Resources, A.B., J.H.P., Y.H., Z.L., C.A.W., L.F., A.B.S., M.R.C.;

Writing – original draft, S.K., Y.A.Q.;

Writing – review & editing, all authors;

Visualization, S.K., Z.L., C.A.W., C.T.;

Supervision, A.B.S., M.R.C., Y.A.Q.;

## Graphical Abstract

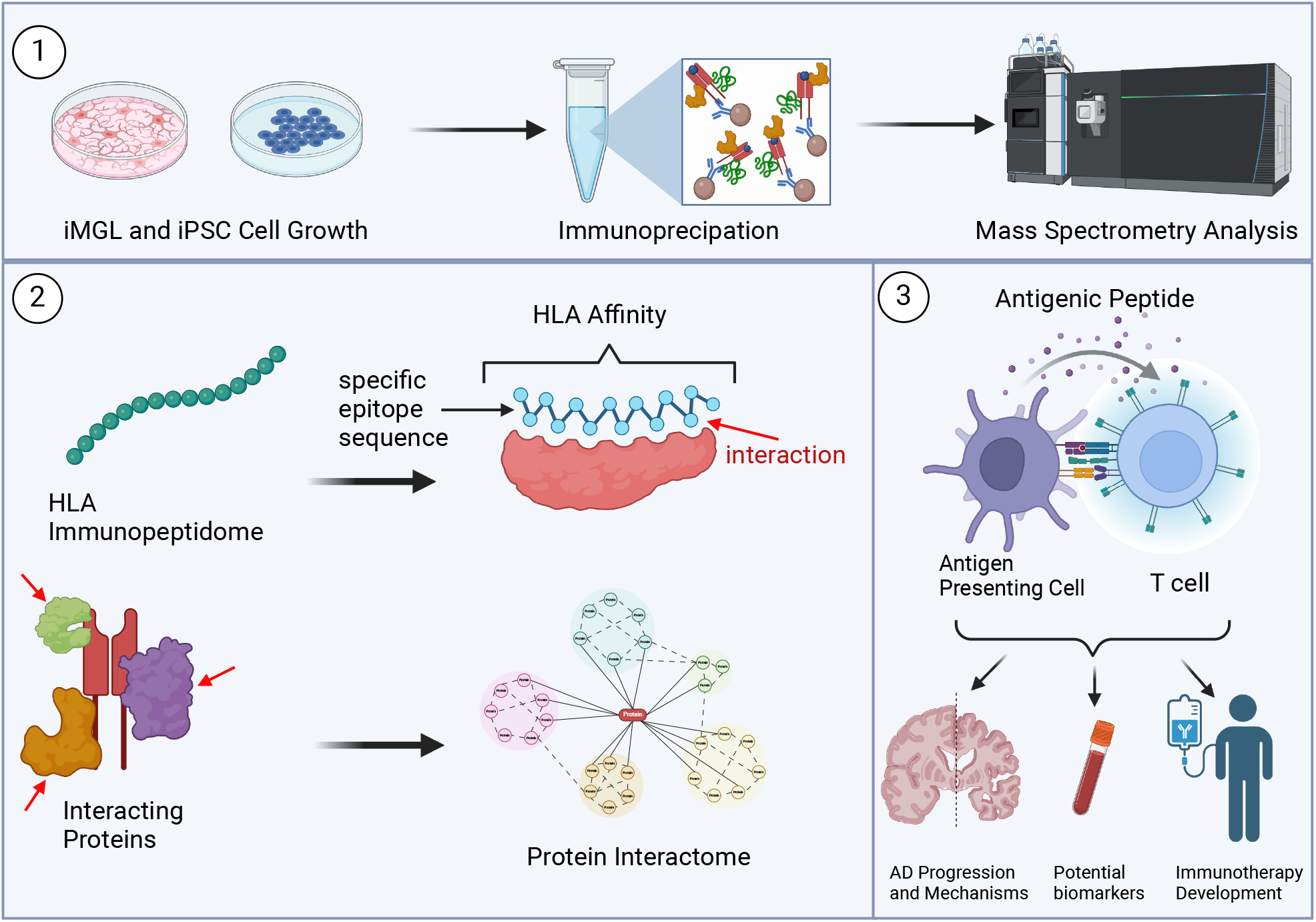

