## Supplemental material for "Large-scale HLA immunopeptidome and interactome profiling in microglia"

**Supplementary Figure 1. Antigen presentation process in brain glia cells.** Microglia process and present foreign epitopes from aggregated proteins that are hallmark in neurodegenerative diseases such as tau and amyloid beta. This presentation causes the recruitment of CD8+ or CD4+ T cells and they will initiate an immune response leading to neuronal damage and neuroinflammation as observed in many neurodegenerative diseases.

**Supplementary Figure 2. Total proteome of HLA expression in sample types.** Box plot showing the HLA-I and HLA-II abundance in iMGL and iPSCs. Statistical analysis was performed using a two-way ANOVA with Tukey's multiple comparisons test (**** $p < 0.0001$, *** $p < 0.001$, ** $p < 0.01$, * $p < 0.05$).

**Supplementary Figure 3. HLA immunopeptidome deconvolution. (A)** PCA plot showing peptidome variance between iMGL and iMGL with IFNγ-treatment. PC1 presented with 27.52% variance and PC2 presented with 15.31% variance. **(B)** HLA-I peptide affinity count plot that displays number of peptides from data that interact with an HLA-II allele. NetMHCpan version 4.3 was used for affinity prediction and cutoff rank was set to < 2%. **(C)** HLA-II peptide affinity count plot that displays number of peptides from data that interact with an HLA-II allele. NetMHCpan version 4.3 was used for affinity prediction and cutoff rank was set to < 5%.

**Supplementary Figure 4. Pathway analysis of unique HLA PPIs in iPSCs or iMGL. (A-B)** GO biological processes pathways of unique HLA PPIs to iPSCs **(A)** and iMGL **(B)**. **(C-D)** GO biological processes pathways of unique PPIs to HLA-I **(C)** and HLA-II **(D)** in iPSCs. **(E-F)** GO biological processes pathways of unique PPIs to HLA-I **(E)** and HLA-II **(F)** in iMGLs.

**Supplementary Figure 5. Pathway analysis of activated and suppressed HLA PPIs in iMGL compared to iPSCs. (A-B)** GO pathway plot **(A)** and KEGG pathway **(B)** for significantly altered HLA-I interacting proteins in iMGL compared to iPSCs. Color intensity dictates adjust p-value; Bubble size represents the amount of the corresponding interaction proteins in each pathway. The GeneRatio is calculated by (count of core enrichment genes) / (count of pathway genes). **(C-D)** GO pathway plot **(C)** and KEGG pathway **(D)** for significantly altered HLA-II interacting proteins in iMGL compared to iPSCs. Color intensity dictates adjust p-value; Bubble size represents the amount of the corresponding interaction proteins in each pathway. The GeneRatio is calculated by (count of core enrichment genes) / (count of pathway genes).

**Supplementary Figure 5. HLA interactome network.** STRING diagram of proteins that are involved and interact with the HLA protein and its mechanism. In GO pathway 0002483 (Antigen processing and presentation of endogenous peptide antigen), 11 interacting proteins were identified and presented with an FDR of 1.25 E-28, with a confidence of 0.9. In GO pathway 0002478 (Antigen processing and presentation of exogenous peptide antigen), 15 interacting proteins were identified and presented with an FDR of 3.69 E-32, with a confidence of 0.9.

**Supplemental Table S1.** Intensity of whole-cell proteins identified in iPSC, iMGLs, iMGLs treated with IFNγ

**Supplemental Table S2.** Intensity of HLA peptide identified in iPSC, iMGLs, iMGLs treated with IFNγ

**Supplemental Table S3.** Risk genes and variants associated with Alzheimer's disease and related dementias

**Supplemental Table S4.** Intensity of HLA interaction proteins identified in iPSC, iMGLs, iMGLs treated with IFNγ
