## Supplemental figures for "Large-scale HLA immunopeptidome and interactome profiling in microglia"

### Supplementary Figure 1

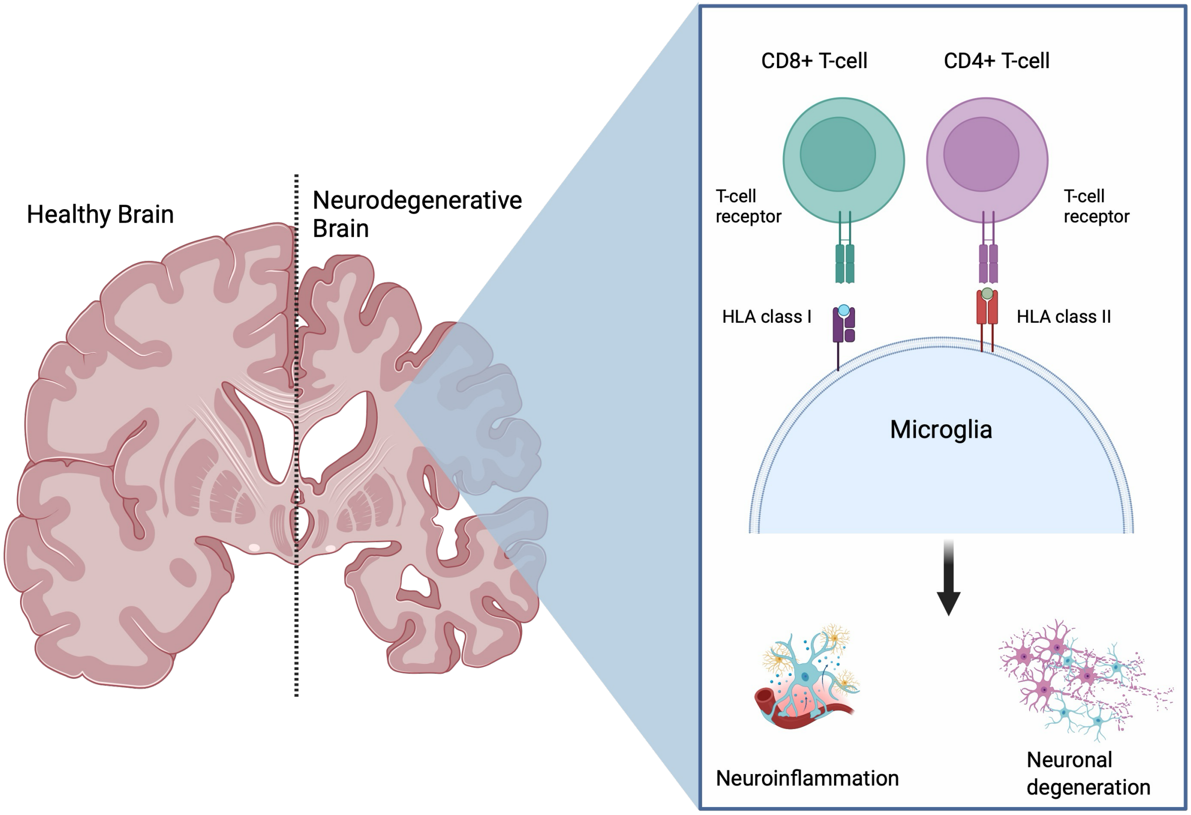

**Supplementary Figure 2**

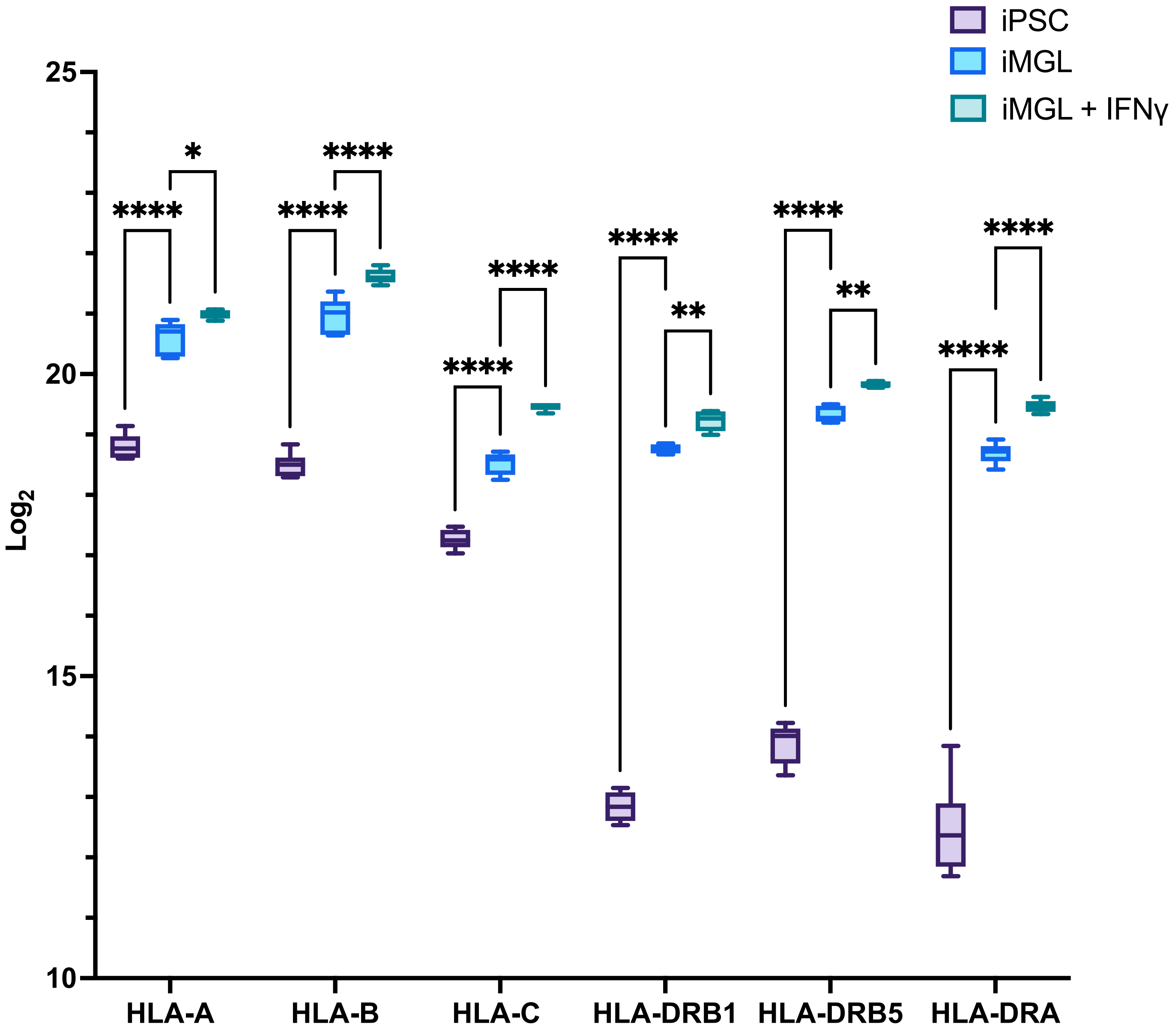

### Supplementary Figure 3

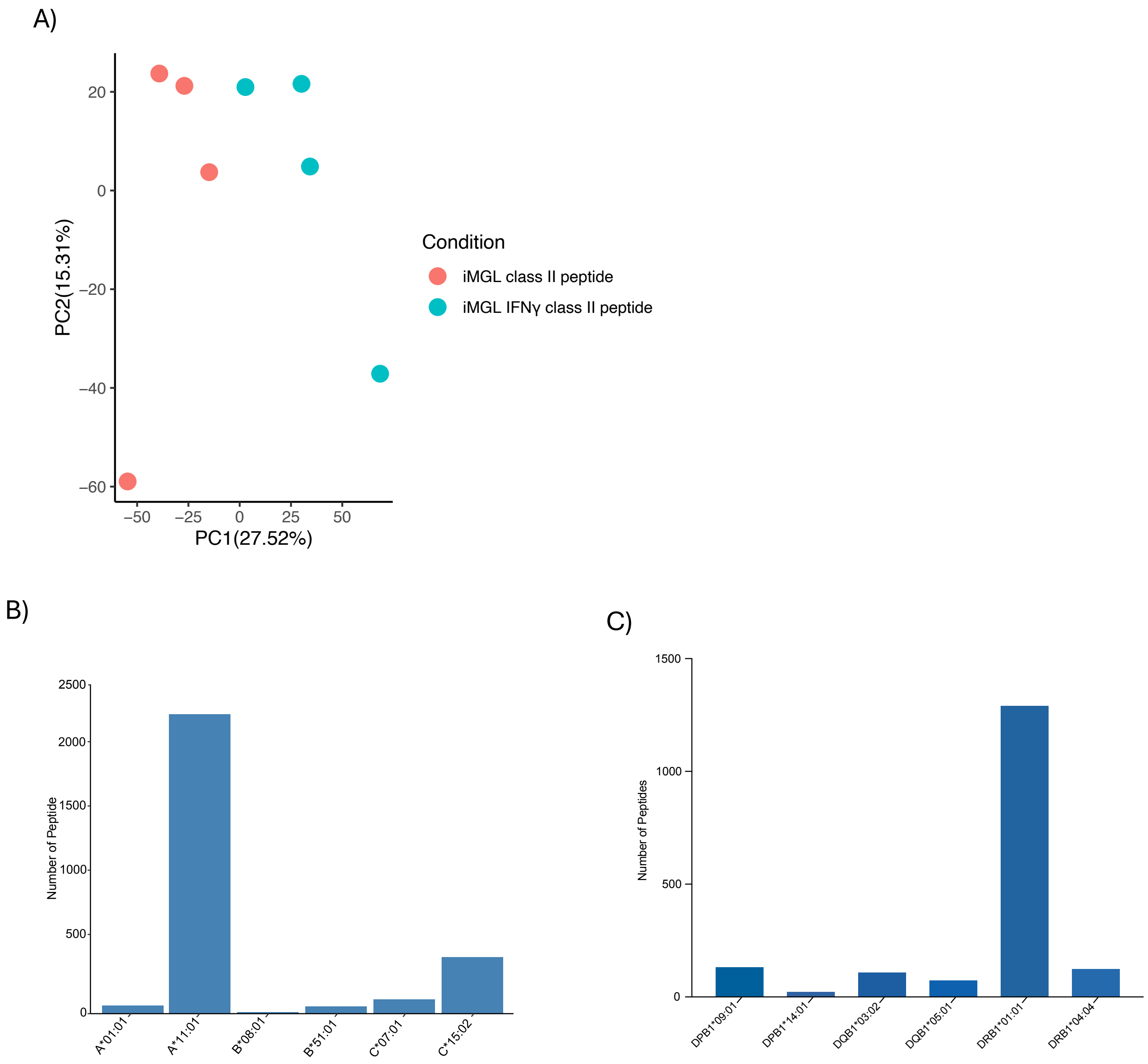

### Supplementary Figure 4

A)

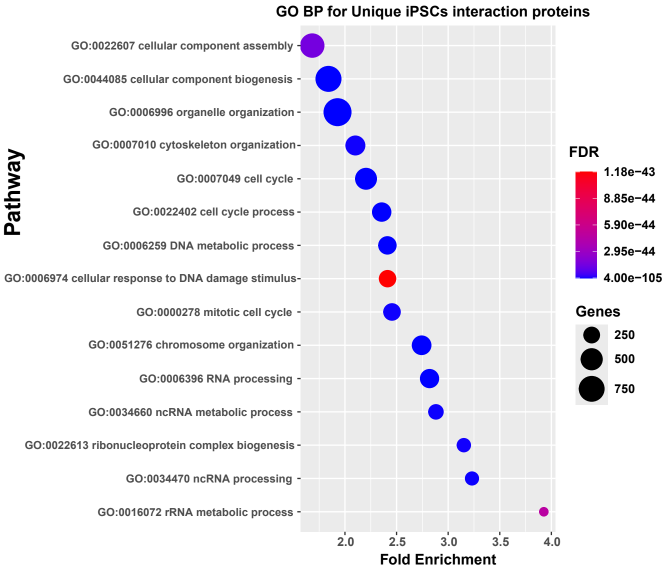


B)

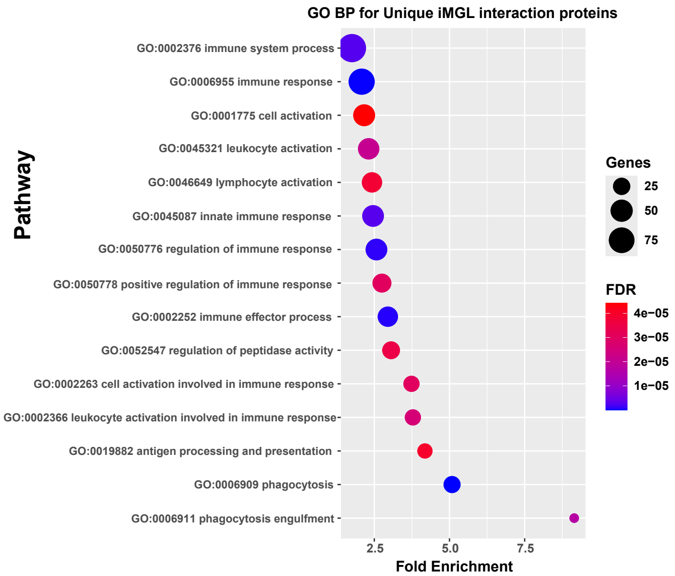


C)

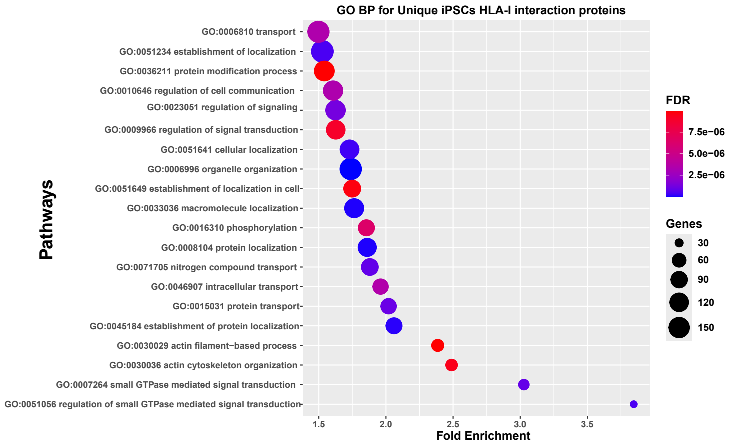


D)

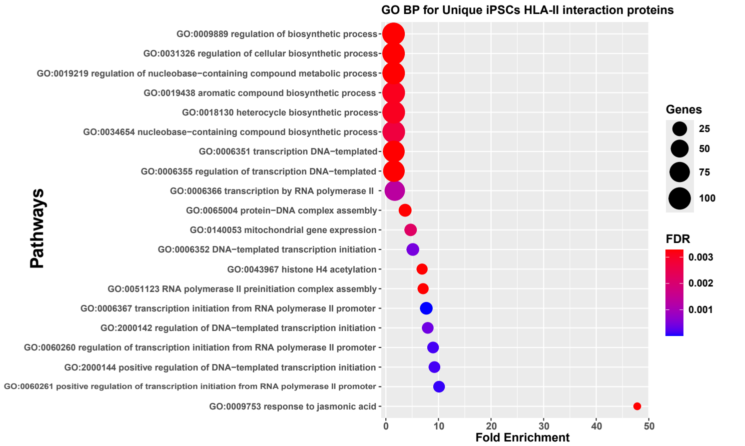


E)

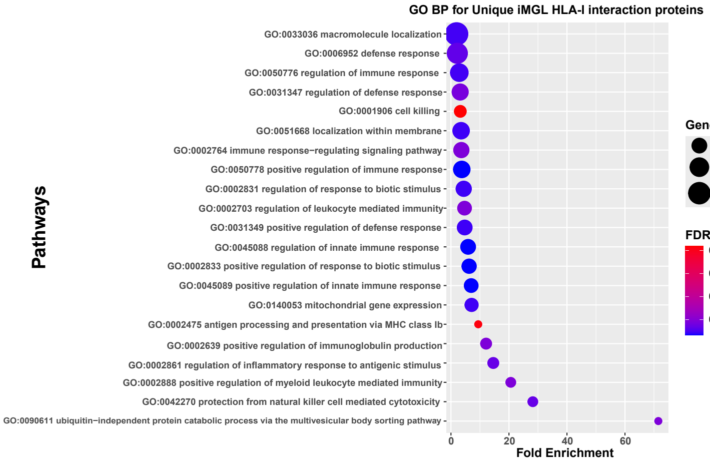


F)

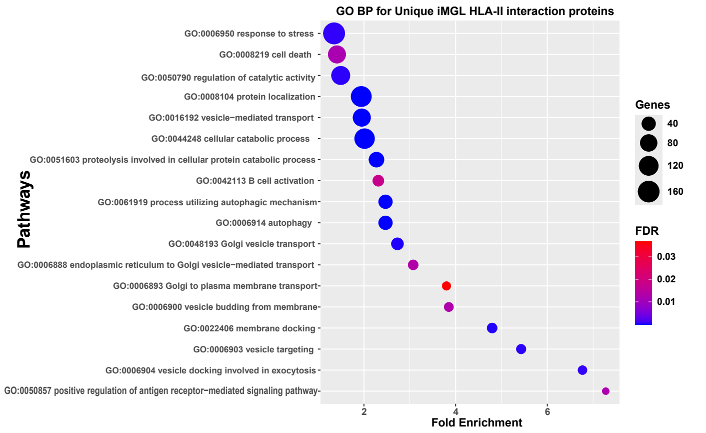

### Supplementary Figure 5

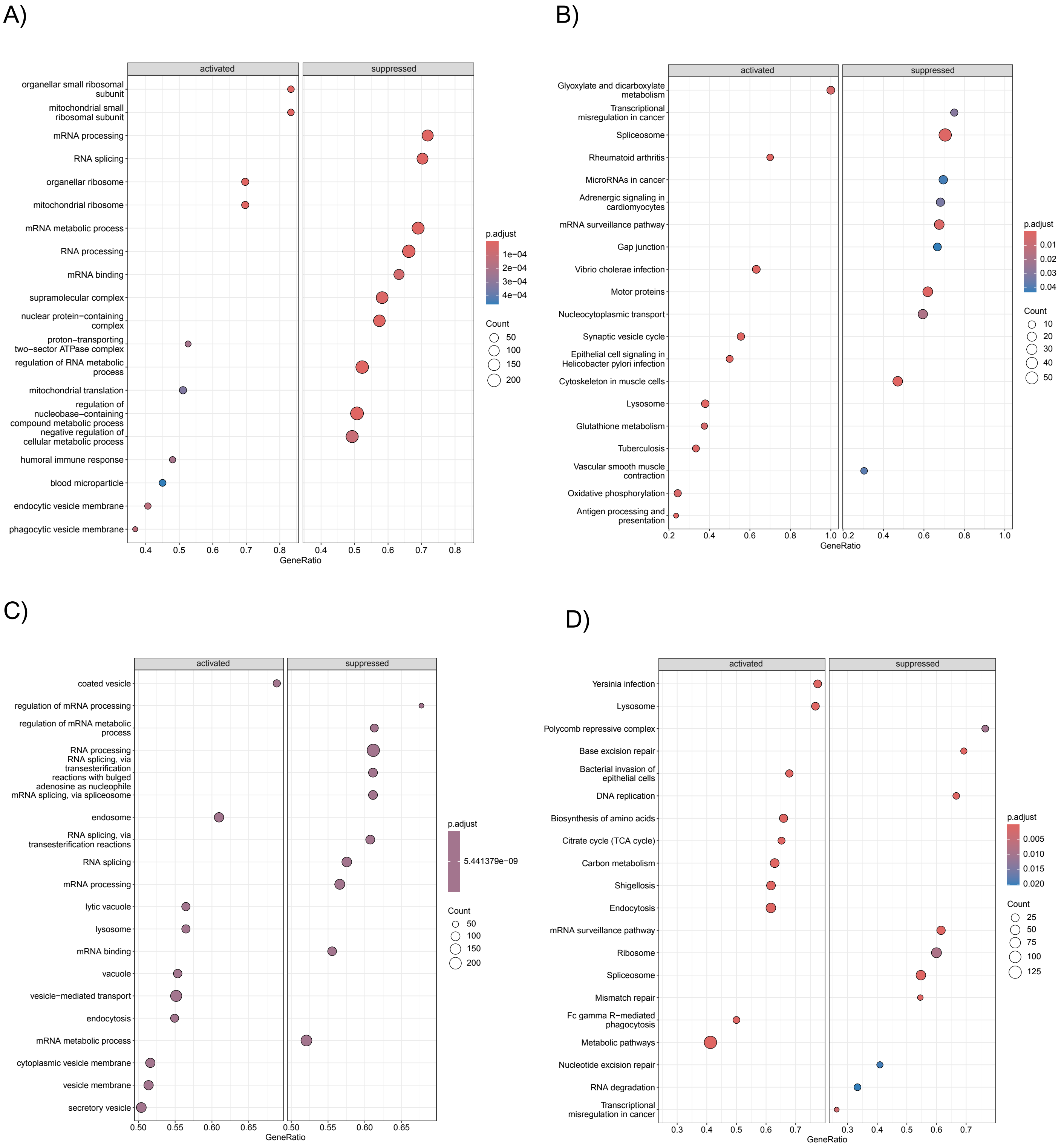

**Supplementary Figure 6**

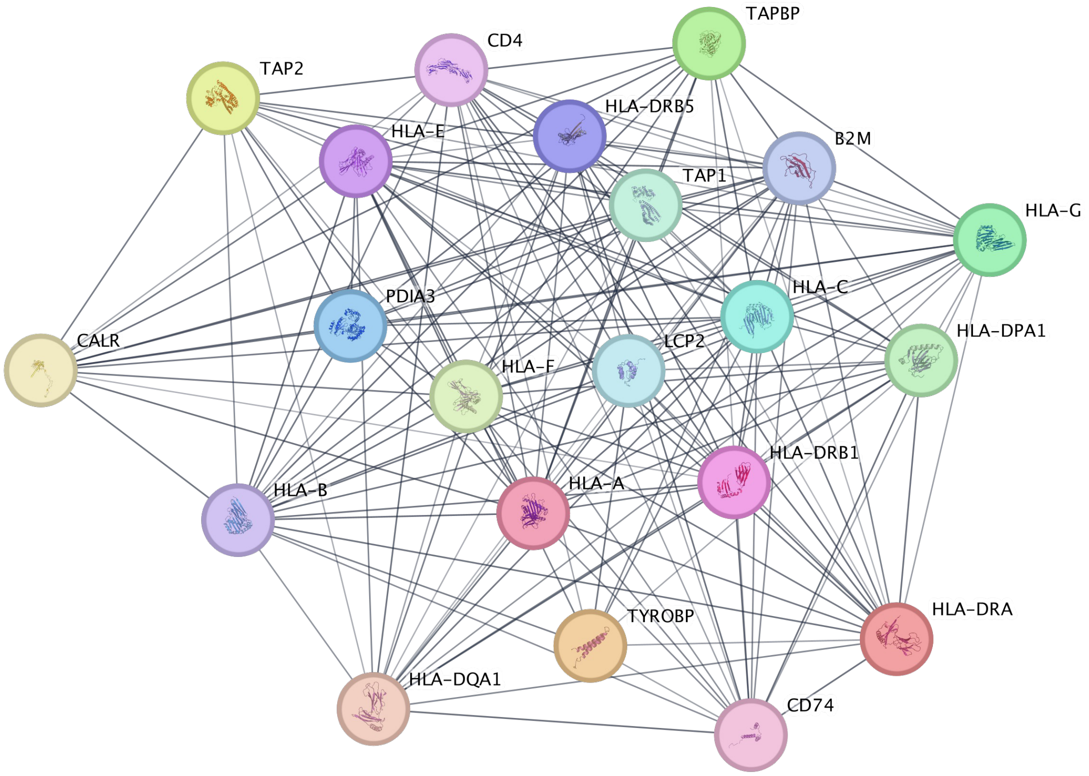
